# Lipid nanoparticle topology regulates endosomal escape and delivery of RNA to the cytoplasm

**DOI:** 10.1101/2022.05.20.492895

**Authors:** Lining Zheng, Sarith R. Bandara, Cecilia Leal

## Abstract

RNA therapeutics have the potential to resolve a myriad of diseases caused by gene deficiency. Lipid nanoparticles (LNPs) are one of the most successful RNA delivery systems. However, expanding their application hinges on the discovery of next generation LNPs with high potency, cyto-specific targeting, and low side effects. Overcoming the difficulty of releasing cargo from endocytosed LNPs remains a significant hurdle. The endosomal escape of viral and non-viral nanoparticles relies on the topological transformation of membrane fusion pore formation followed by RNA translocation into the cytosol. In this study we show that LNP-RNA nanostructure modulates the energetic cost of LNP fusion with a target membrane. The inclusion of a new class of structurally-active lipids leads to superior LNP endosomal fusion, fast evasion of endosomal entrapment, and efficacious RNA delivery. Specifically, bicontinuous cubic RNA-LNPs, ***cuboplexes***, have significantly higher endosomal escape rates and deliver more RNA compared to regular lamellar LNPs.

## 1 Main

RNA therapeutics has gained wide attention due to its power to resolve insufficient expression of critical proteins that drive a myriad of human diseases including cancer [1], neurodegeneration [2], metabolic disorders [3], among others[4][5][6] [7][8]. Delivery vectors play an important role in the development of such therapeutics in preventing RNA degradation and successfully deliver RNA to target cells. Compared to viral vectors, non-viral systems have the advantage of being less immunogenic and easier to manufacture [9]. Among the different materials used for non-viral delivery, lipid nanoparticles (LNPs) have been particularly successful, as many LNP-RNA formulations are clinically available[10][11]or have advanced to clinical trials. The first FDA approved small interfering RNA (siRNA) treatment, patisiran[12], utilized LNPs for delivery. The effectiveness of LNPs has been further manifested by the development of lipid-mRNA (messenger RNA) vaccines for coronavirus disease 2019 (COVID-19) [13].

Despite the astonishing progress LNP delivery achieved, many barriers are still present in realizing the full potential of LNP-RNA systems[14]. While most LNP systems are efficiently taken up by the cell via endocytosis, they often remain trapped in endosomal compartments and degrade through the endosome-lysosome acidification pathway [15] [16] [17] [18]. One of the main bottlenecks of LNP-based RNA delivery is poor endosomal escape[19] [20]. Viral delivery systems have membrane proteins that undergo a conformation change allowing them to easily fuse with the plasma or endosomal membranes to release their payload. Previous studies have shown that for LNP mediated siRNA delivery, less than 2% of the siRNA can successfully escape the endosome and reach the cytosol [21]. With low cytosolic delivery efficacy, much research has been done in attempt to improve endosomal escape and transfection efficiency of LNPs by modifying lipid compositions[22][23][24]. Sahay and coworkers have shown that LNPs containing cholesterol demonstrate improved delivery efficiencies, potentially caused by enhanced endosomal membrane fusion [25]. Other ways of optimizing lipid compositions also shown success [26], such as the inclusion of ionizable lipids [27][28] [29]. Ionizable lipids are able to stay neutral under physiological pH but become cationic under the acidic environment of endosomes, being able to electrostatically pin the endosomal membrane which may allow LNPs to destabilize it and facilitate escape [10].

However, compared to the extensive research done exploring different lipid formulations, the nanostructure of LNPs, or the specific packing of lipids and nucleic acids into the LNP has not been fully investigated. In this paper we demonstrate that future LNP design should include a new class of structurally active lipids. Due to their amphiphilic nature, lipid molecules can form a variety of self-assembled structures in aqueous environments (Fig.1b) that are conserved when encapsulating nucleic acids. This includes lamellar vesicles (*L*), inverse hexagonal (*H*_*II*_) and bicontinuous cubic phases (*Q*_*II*_) [30][31][32][33][34][35][36][37][38][39][40]. The optimal mechanism to boost LNPs endosomal escape is to promote LNP-endosomal membrane fusion, a process that should take place in less than 30 seconds [41] and be independent from endosome acidification or the debated “proton sponge” effect [42][43]. The design of “fusogenic” LNPs has been exclusively attributed to tuning the molecular packing parameter of lipid molecules such that their spontaneous membrane curvature (*C*_0_) is negative [44][45]. However, the elastic energy cost of membrane fusion or, more importantly, the development of the required fusion pore is controlled by a topological transformation that should mostly depend on the Gaussian moduli of membranes 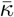. The membrane elastic energy can be represented by the Helfrich equation [46][47], 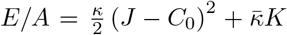, where *κ* is the bending modulus, *J* the total/extrinsic curvature which equals the sum of the principle curvatures *C*_1_ + *C*_2_, and *C*_0_ is the spontaneous curvature. The second term comprises the Gaussian curvature *K* = *C*_1_*C*_2_ and the Gaussian modulus 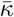. Taking into account the Gauss-Bonnet theorem [48] and the Helfrich framework, one can calculate that the elastic energy cost of the topological transformation from two nested membrane vesicles 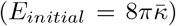 to two fused vesicles 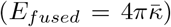 to be 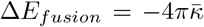 [49] (Fig.1a). It is immediately clear that modulating the Gaussian modulus 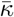 and Gaussian curvature *K* will significantly impact membrane fusion events and the formation of fusion pores through which RNA cargo can be delivered into the cytosol (Fig.1c). Of the main categories of reported LNP nanostructures (*L, H*_*II*_, and *Q*_*II*_), positively curved vesicular lamellar phases (*L*) have 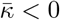 but cubosomes [50] or cuboplexes (cubosomes loaded with RNA [35][36]) which are made of bicontinuous cubic phases have intrinsically negative Gaussian curvature *K* and positive 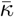 and the activation energy for fusion with endosomal membranes and fusion pore formation should be the lowest. D. P. Siegle [51][52][53][54][55][56] first suggested that bicontinuous cubic phases should be highly fusogenic and we conjecture that *Q*_*II*_ LNPs have a greater potential in fusing with the endosomal membrane and successfully release their cargo (Fig.1c).

In this paper we show that the structure of LNP-RNA complexes plays a dominant role in the extent of endosomal-membrane fusion and we enable new design principles for the development of highly efficient LNP-based RNA delivery systems.

**Fig. 1.**
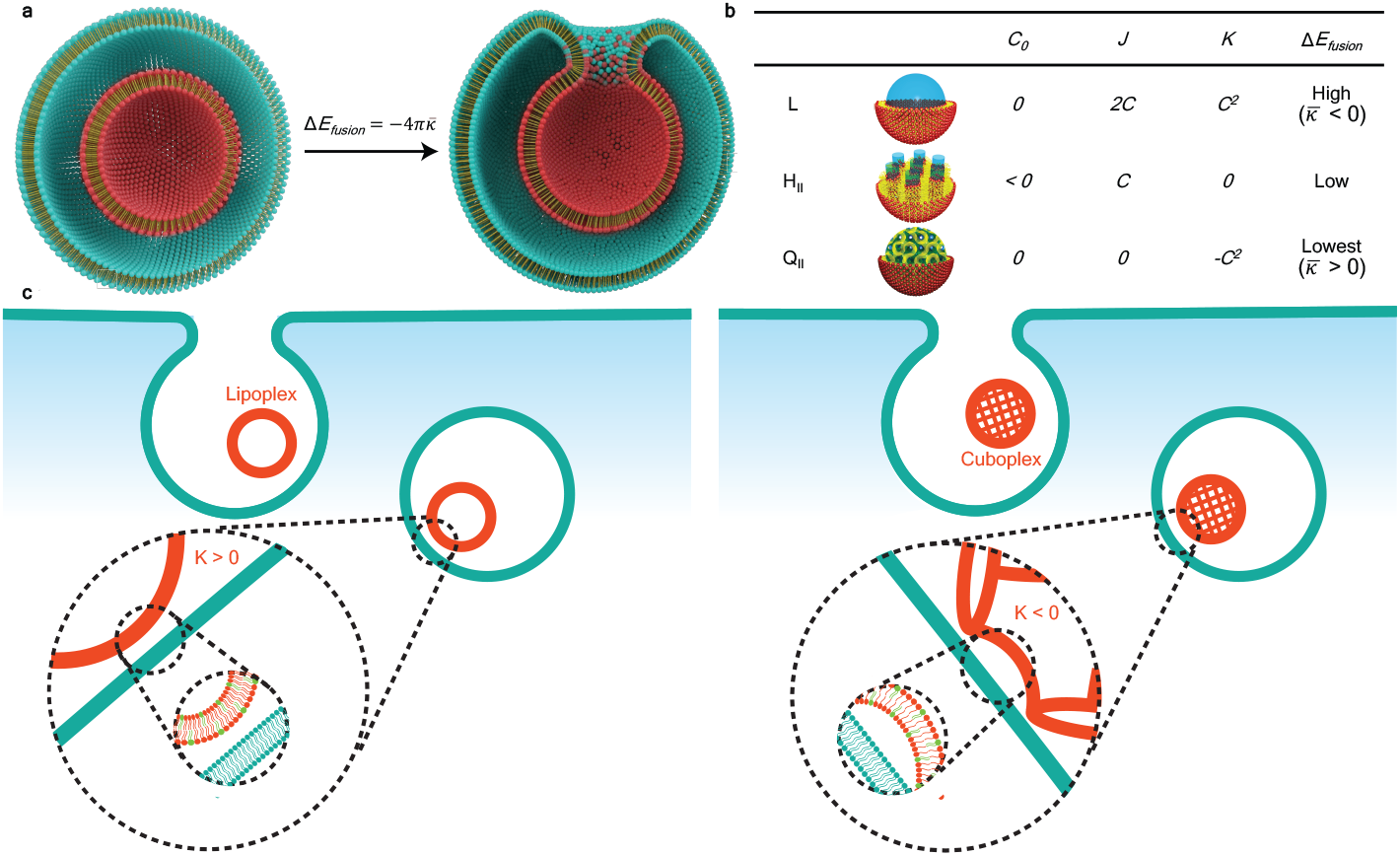
Schematic representation of the cell entry of LNP-RNA complexes. **a**, LNPs (red particle) enter the cells via endocytosis and are entrapped in the endosome (teal vesicle). Illustration credits: Alex D. Jerez, Imaging Technology Group at the Beckman Institute University of Illinois, Urbana-Champaign. LNP-endosome membrane fusion involves a topological transformation of activation elastic energy 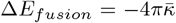. **b**, LNPs of different nanostructures (*L, H*_*II*_, and *Q*_*II*_) have distinct spontaneous curvature *C*_0_, total curvature *J*, and Gaussian curvature *K* properties. **c**, The escape of LNPs from endosomal entrapment will be enhanced by lowering the energetic barrier for fusion and fusion pore formation with the endosomal membrane for which *Q*_*II*_ LNPs should have preferential properties.

### 1.1 Formation of LNP-RNA complexes of precise nanostructures

In our study, we judiciously select specific neutral lipid molecular systems and compositions to control the nanostructure of LNP-RNA complexes. Glycerol monooleate (GMO) is a neutral lipid approved by the Food and Drug Administration (FDA) for *in vivo* use and is utilized as an adjuvant [57][58]. GMO is well-known to be stable into a variety of bicontinuous cubic phases [59] that be formulated into cubosome LNPs (*Q*_*II*_) encapsulating siRNA which we termed as cuboplexes [35][36]. The cuboplex LNP formulation comprised GMO, a cationic lipid 1,2-dioleoyl-3-trimethylammonium propane (DOTAP) just enough to electrostatic pin RNA without raising toxicity, as well as 1,2-dioleoyl-sn-glycero-3-phosphoethanolamine-N-[methoxy(polyethylene glycol)-2000 (DOPE-PEG) to optimize the colloidal stability of the LNPs [60] [61] [62]. Modulating the amount of neutral lipid GMO in the ternary GMO, DOTAP, and DOPE-PEG mixture results into precise tuning of different bicontinuous cubic structures of distinct space groups (*Ia*3*d* - gyroid, *Im*3*m* - primitive, and *Pn*3*m* - diamond) as well as the formation of hexagonal phases (*H*_*II*_) [35][30]. To form the traditional lamellar phase (*L*), GMO contents are low (*<*25 mol%) [30] or the neutral lipid was switched to a phosphatidylcholine 1,2-dioleoyl-sn-glycero-3-phosphocholine (DOPC). LNP-RNA complexes are formed with an optimal [35] charge ratio of *ρ*=3, where charge ratio represents the number of positive charges from lipid molecules (*n*_*DOTAP*_) over the number of negative charges from RNA (*n*_*NA*_). When nucleic acid is added to the lipid systems the nanostructures are mostly conserved but there is a preference to adopt *H*_*II*_ phases as the 1D straight cylindrical water channels can accommodate long chain nucleic acids [63]. Fig. 2 shows structural information of dilute LNP-RNA complexes (siRNA and mRNA) at different amounts of neutral GMO obtained by synchrotron small-angle X-ray scattering (SAXS). LNP-siRNA complexes with a GMO/DOTAP/DOPE-PEG (molar ratio 85/14/1) composition show four distinct diffraction peaks that indicate the presence of a bicontinuous cubic gyroid nanostructure (*Q*_*II*_) coexisting with a 2D inverse hexagonal phase (*H*_*II*_) (Fig. 2a). The first two peaks showr eciprocal lattice vectors 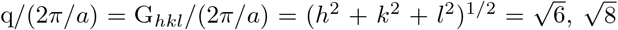, which corresponds to the {211}, {220} planes of a Q_*II*_ gyroid phase. The first, third and fourth peaks show reciprocal lattice vectors 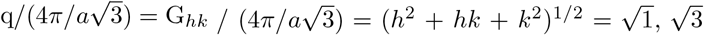, and 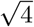, corresponding to {10},{11}, and{20} of the *H*_*II*_ phase. The lattice spacings are 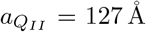 and 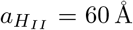. LNP-mRNA complexes with the same lipid composition have a similar scattering profile and display *H*_*II*_ / *Q*_*II*_ phase coexistence with lattice spacings 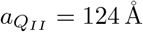 and 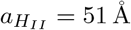 At intermediate GMO contents (Fig. 2b) LNP-RNA complexes mostly adopt the *H*_*II*_ phase in coexistence with the *L* phase. The lattice spacings of the complexes are similar, with LNP-siRNA 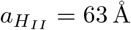 and *a*_*L*_ = 59 Å and LNP-mRNA 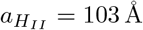 and *a*_*L*_ = 91 Å. At low GMO content (below 25 mol%) LNP-RNA complexes adopt the *L* phase (Supplementary Fig. 1). Switching GMO for a phosphatidylcholine neutral lipid always results in a *L* phase shown in Fig. 2c for DOPC/DOTAP/DOPE-PEG (molar ratio 85/14/1). The complexes show similar lattice parameters, with LNP-siRNA *a*_*L*_ = 332 Å and LNP-mRNA 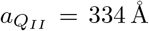The modulation of neutral lipid amount and identity yields rich structural diversity of LNP-RNA complexes without disrupting their stability. LNP-RNA complexes remain colloidally stable regardless of nanostructure having size distributions between 150 and 250 nm (Supplementary Table 1).

**Fig. 2.**
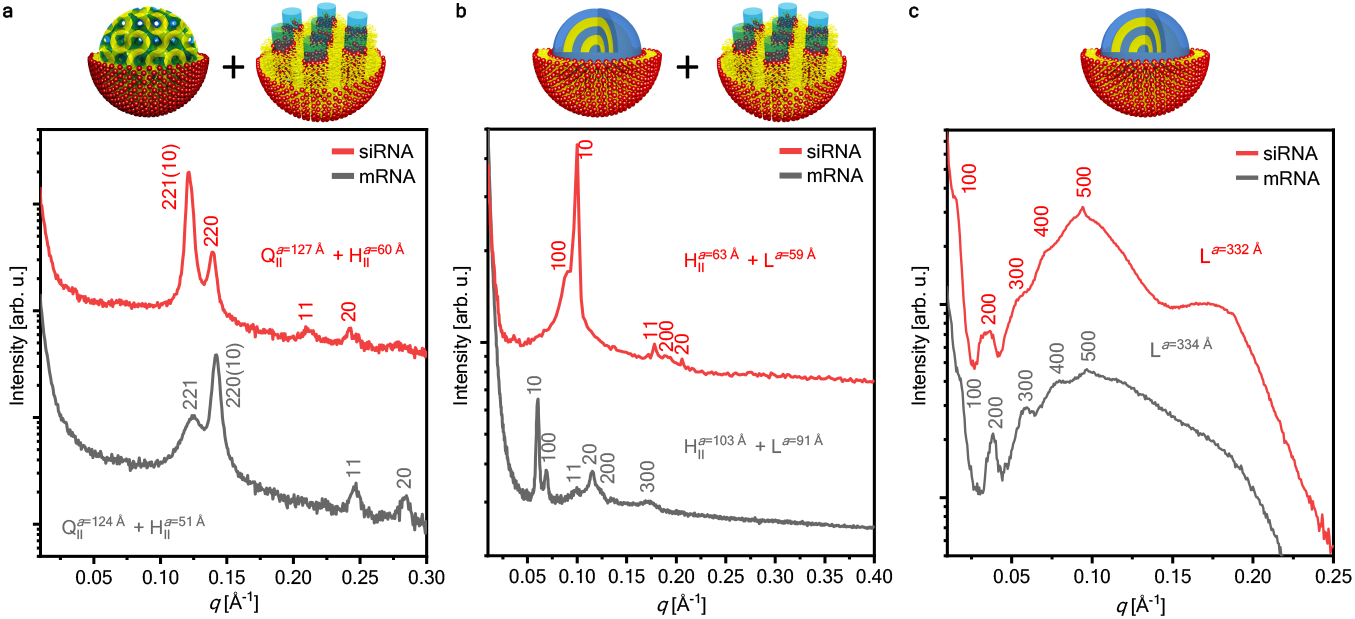
Synchrotron SAXS data of LNP-siRNA and LNP-mRNA complexes under dilute conditions show bicontinuous cubic (*Q*_*II*_), 2D reversed hexagonal (*H*_*II*_) and lamellar phases (*L*). **a**, *Q*_*II*_, *H*_*II*_ LNP-RNA complexes were formed with a lipid composition of GMO/DOTAP/DOPE-PEG at molar ratio 85/14/1 and mixed with siRNA or mRNA at *ρ*=3. **b**, *H*_*II*_, *L* LNP-RNA complexes were formed with a lipid composition of GMO/DOPC/DOTAP/DOPE-PEG at molar ratio 50/35/14/1 and mixed with siRNA at *ρ*=3. **c**, *L* LNP-RNA complexes were formed with a lipid composition of DOPC/DOTAP/DOPE-PEG at molar ratio 85/14/1 and mixed with siRNA or mRNA at *ρ*=3.

### 1.2 Fusion of LNP-RNA complexes with endosomes depends on LNP nanostructure

The interaction between endosomes and LNP-RNA complexes of different nanostructures *in* − *vitro* were explored using confocal laser scanning microscopy (CLSM), fluorescence resonance energy transfer (FRET) assays, and live-cell self-quenching CLSM experiments (Fig. 3). Fig. 3a shows CLSM images of isolated endosomes incubated with pure LNPs (no RNA) having the *L* (liposome) and *Q*_*II*_ cubosomes nanostructures. Endosomes were isolated by standard spin–column techniques from HeLa cells and fluorescently labeled by incubating in 20*µ*M Nile Red. The LNPs were tagged with 0.1% 1,1’-dioctadecyl-3,3,3’,3’-tetramethylindodicarbocyanine (DiD). LNPs do not aggregate significantly on their own but when incubated with endosomes, LNP-endosome fusion and aggregation is readily visible by co-localization of DiD and Nile Red fluorescence in both systems but is significantly more pronounced for *Q*_*II*_ LNPs.

**Fig. 3.**
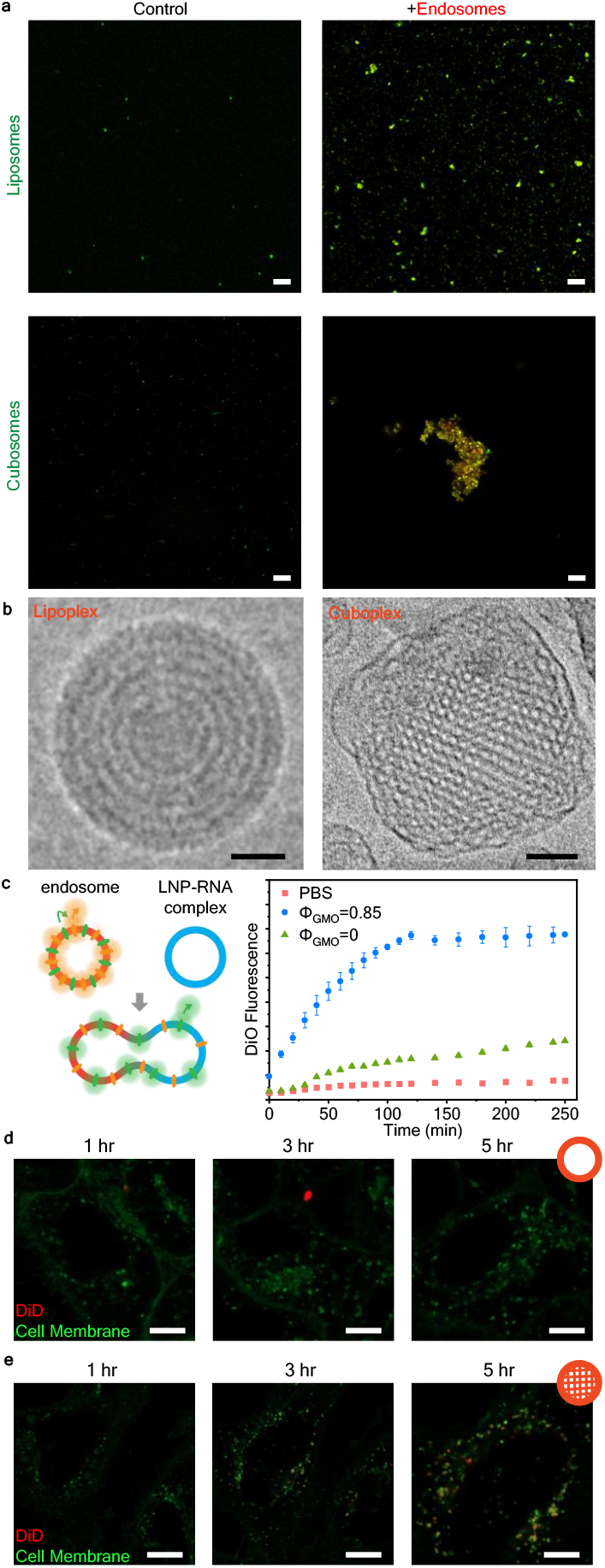
Cubic and hexagonal LNPs fuse more readily with endosomes compared to lamellar LNPs. **a**, Visualization of LNP (green) and endosome (red) interaction with CLSM after 6 hr incubation. Scale bar, 20 *µ*m. **b**, Representative Cryo-EM images of lipoplexes and cuboplexes. Scale bar, 50 nm. Reproduced with permission from ref [36]. Copyright 2018 American Chemical Society. **c**, FRET assay to evaluate membrane fusion between endosomes and LNP-siRNA complexes of different structures. The fusion extent is indicated by DiO fluorescence recovery (*n* = 3, data presented as mean ± s.d.).**d-e**, Live cell imaging of HeLa cells treated with lipoplexes (**d**) and cuboplexes (**e**) labeled with 1% self quenching dye DiD. Recovery of DiD signal (red) implies LNP-endosomal fusion. Scale bar, 10 *µ*m.

After observing that *Q*_*II*_ LNPs bind with isolated endosomes more favorably, we conducted a FRET assay[64] to evaluate the extent of the fusion process between endosomes and RNA–loaded LNPs of different nanostructures. Isolated endosomes were co–labeled with 1,1’-dioctadecyl-3,3,3’,3’-tetramethylindocarbocyanine perchlorate (DiI) and 3,3’-dioctadecyloxacarbocyanine perchlorate (DiO), with DiO acting as a donor fluorophore and DiI acting as an acceptor fluorophore. The proximity of the dye molecules enables FRET to occur leading to enhanced acceptor (DiI) fluorescence. If endosomes fuse with another membrane, the dye molecules diffuse, increasing the distance between each other and FRET occurrences diminish. Reduced FRET allows donor (DiO) fluorescence to recover. Thus, the extent of membrane fusion can be evaluated by the increase in DiO fluorescence. To see how LNP structure influences LNP-endosome membrane fusion, we prepared LNP-siRNA complexes with same DOTAP molar fractions (Φ_*DOTAP*_ = 0.14) but different GMO molar fractions. At Φ_*GMO*_ = 0.85 RNA–loaded LNPs display the *Q*_*II*_ structure (in coexistence with *H*_*II*_) termed cuboplexes[35][36] and at Φ_*GMO*_ = 0 the lamellar *L* phase (Fig. 2) termed lipoplexes. Fig. 3b shows Cryogenic electron microscopy (Cryo-EM) images of cuboplexes and lipoplexes LNPs[36]. The intensity of DiO fluorescence as a function of time (Fig. 3c) is significantly higher for LNPs comprising Φ_*GMO*_ = 0.85 (blue data points) compared to Φ_*GMO*_ = 0 (green data points). This is consistent with a considerably higher extent of fusion of endosomes with ′′non-lamellar′′ LNPs (*Q*_*II*_*/H*_*II*_ compared to regular “lipoplex” LNPs. Additional data is shown in Supplementary Fig. 2..

The fact *Q*_*II*_ RNA–loaded LNPs fuse with isolated endosomes side-by-side on a glass slide is encouraging for their application as fusogenic LNPs for RNA delivery but this is not representative of the fusion process in live cells where LNPs have to break out of an enclosing endosomal membrane. To evaluate how LNPs of different nanostructures fuse with the entrapping endosomal membranes in live cells we conducted a fluorescence self–quenching experiment in live HeLa cells (Fig. 3d,e). We employed two types of siRNA-loaded LNPs: cuboplexes (*Q*_*II*_) and lipoplexes (*L*)) labeled with high molar percentages (1%) of a self-quenching dye, DiD [65] [66]. When the membranes of the LNP-siRNA complexes fuse with endosomes, DiD molecules will diffuse across the membrane, reducing the self-quenching of DiD and increasing its fluorescence. Fig. 3d,e displays the obtained DiD fluorescence recovery at three time points (1, 3, and 5 *h*) after incubation with *L* (Fig. 3d, lipoplexes) and *Q*_*II*_ (Fig. 3e, cuboplexes) siRNA loaded LNPs. As time progresses, an increasing DiD signal (red fluorescence signal) is observed in cells treated with cuboplexes, indicating that indeed LNPs successfully fused with the endosomal membrane that internalized them. Comparatively at the same time point, cells treated with lipoplexes show little DiD fluorescence recovery, implying less effective endosomal membrane fusion. These results show that the nanostructure of LNP-RNA complexes plays a critical role in their ability to fuse and break out of endosomal compartments.

### 1.3 RNA delivery of LNP-RNA complexes of different nanostructures

We next investigated if the ability for *Q*_*II*_ RNA-loaded LNPs (cuboplexes) to efficiently fuse with endosomal membranes translates to better delivery of cargo and more efficient endosomal escape. Uptake of siRNA in epithelial human cervix cancer (HeLa) and murine breast cancer (4T1) cells were quantified by flow cytometry (Fig. 4a). To verify that the siRNA signal is not from siRNA outside the cell membrane or from dead cells, trypan Blue (TB) was added to quench the fluorescence of siRNA outside the cellular membrane and that of membrane compromised cells [67] [68]. The results show that HeLa cells treated with cuboplexes have significantly more cytoplasmic siRNA compared to cells treated with lipoplexes, and the siRNA uptake is comparable to that of Lipofectamine (LFA), a commercially available siRNA transfection agent. Hela cells are well-known to be relatively easy to transfect, however, in more resilient cells lines like 4T1 murine breast cancer epithelial cells [69], we observe that cuboplexes continue to outperform lipoplexes in siRNA delivery capabilities.

**Fig. 4.**
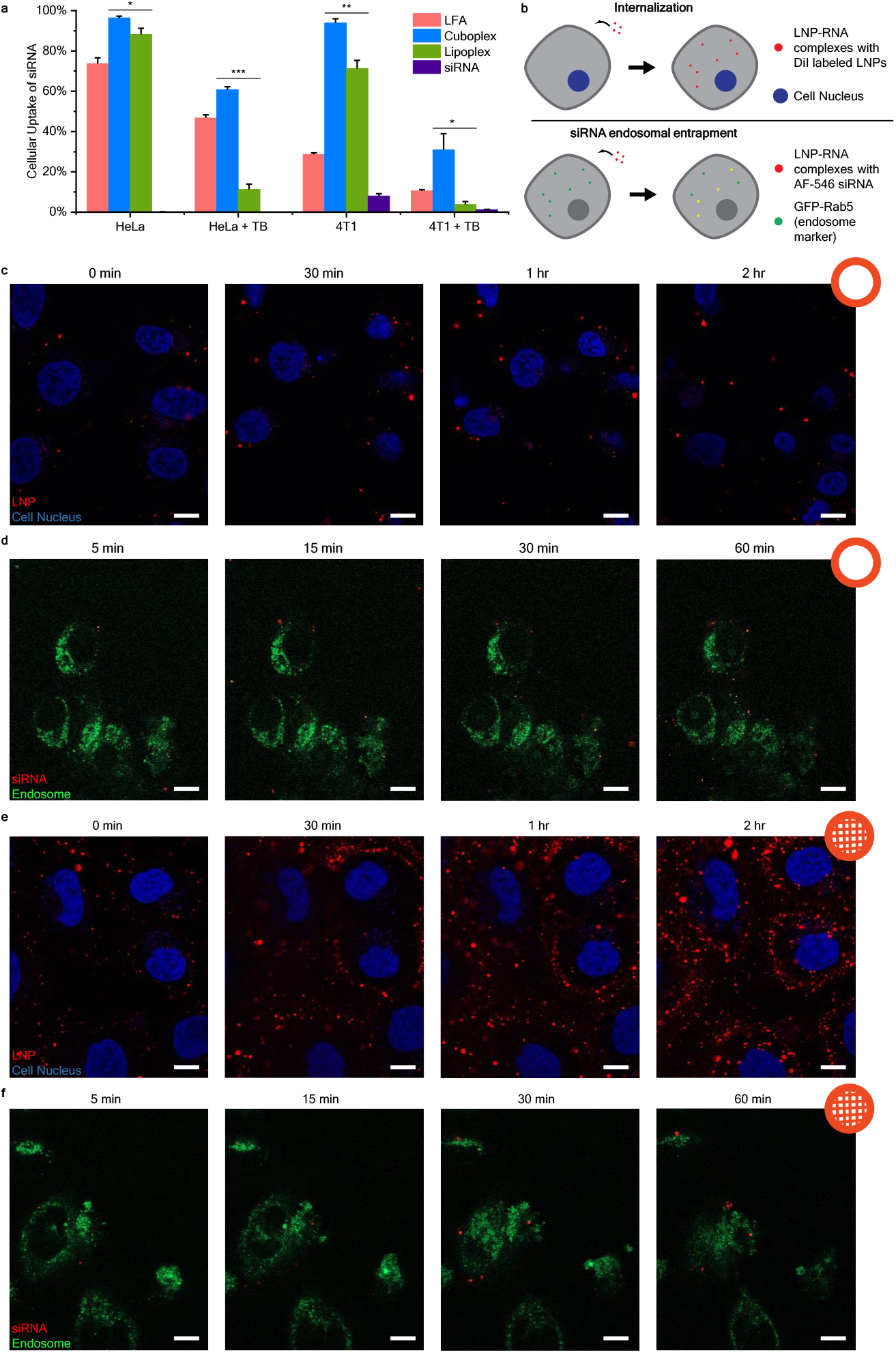
Cuboplexes deliver siRNA more effectively compared to lipoplexes. **a**, Uptake of siRNA in HeLa cells and 4T1 cells quantified by flow cytometry. Trypan Blue (TB) was added to quench the fluorescence of siRNA outside the cellular membrane and that of membrane compromised cells (*n* = 3, data presented as mean ± s.d.). Significance between groups were calculated with two-sided Student’s *t*-test and indicated as \**P ≤*0.05, \*\**P ≤*0.01 or \*\*\**P ≤* 0.001. **b**, Schematic representation of live cell imaging experiments evaluating the internalization and siRNA endosomal entrapment of LNP-RNA complexes. **c-d**, HeLa cells were treated with DiO labeled LNP-siRNA complexes (red) and imaged every 5 min. Less lipoplexes (**c**) enter the cells compared to cuboplexes (**d**). Scale bar, 10 *µ*m. **e-f**, Endosome labeled cells were treated with cuboplexes (**e**) and lipoplexes (**f**) containing AF546-siRNA and imaged every 5 min. Less siRNA (red) are entrapped in the endosome in cuboplex treated cells compared to those treated with lipoplexes. Scale bar, 10 *µ*m.

To test the hypothesis that cuboplexes deliver more RNA to cells compared to lipoplexes because they more effectively fuse and break out of endosomal entrapment, we utilized live cell imaging to compare the cell internalization as well as endosomal entrapment of these two different LNP-RNA complexes (Fig. 4b). DiO labeled LNP-siRNA complexes (shown as red) were added to HeLa cells stained with Hoeschst nuclei dye (blue) and imaged as a function of time (0, 30, 60, and 120 min). The higher amount of red signal around the nuclei would correspond to a higher extent of nanoparticle internalization (Fig. 4 c,e). Lipoplexes show lower levels of internalization by HeLa cells compared to cuboplexes. To investigate the endosomal entrapment of the different LNP-siRNA complexes, early endosomes of HeLa cells were labeled with a fusion construct of Rab5a-GFP (shown as green). The cells were treated with LNP-siRNA complexes prepared with siRNA fluorescently labeled with Alexa Fluor 546 (AF546), shown as red, and monitored over time (5, 15, 30, and 60 min). When siRNA resides entrapped in the endosome, the fluorescent signal of siRNA and endosome would co-localize, appearing as yellow. Less colocalization, or yellow, would be an indication of less endosomal entrapment and more occurrences of successful endosomal escape. In Fig. 4 d,f, compared to lipoplexes, cuboplexes showed less co-localization with the endosomes, indicating less cuboplexes were entrapped.

Combined, lipoplexes show low levels of internalization and high amounts of siRNA-endosome co-localization (Fig. 4 c,d) throughout the investigated timeline of up to 2h (internalization) and 1h (entrapment). This indicates that cuboplexes have superior ability not only to get internalized by cells at faster rates but also to efficiently evade endosomal entrapment and translocate RNA into the cytosol.

We have previously shown that cuboplex LNPs are more efficient at transfecting siRNA into cells compared to lipoplexes leading to better specific gene-knockdown performance.[37] [35] [36]. When replacing siRNA for mRNA LNPs retain their structural identity (Fig. 2) as well as their shape and size (Supplementary table 1). Fig. 5a,b shows images of nanoparticle tracking analysis (NTA) of cuboplex LNPs loaded with siRNA (a) and mRNA (b). Using cuboplex LNPs to deliver mRNA to cells leads to much more efficient activation of luciferase expression (Fig. 5c) compared to parent lipoplex mRNA-LNPs. A plasma membrane integrity assay confirmed that all LNP-RNA complexes used in this study have negligible cytotoxicity (Supplementary Fig.3).

**Fig. 5.**
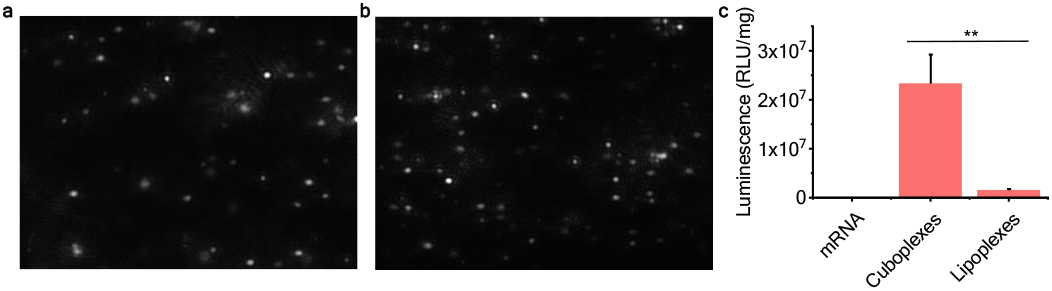
Cuboplexes show higher transfection efficiency of firefly luciferase mRNA compared to lipoplexes. **a-b**, NTA images of cubic LNP-siRNA complexes (**a**) and cubic LNP-mRNA complexes (**b**). **d** Luciferase activity of HeLa cells transfected with firefly luciferase mRNA with mRNA alone, cuboplexes and lipoplexes. (*n* = 3, data presented as mean ± s.d.). Significance between groups were calculated with two-sided Student’s *t*-test and indicated as \**P ≤* 0.05, \*\**P ≤* 0.01 or \*\*\**P ≤* 0.001.

In this study, we were able to show that nanostructures have a significant impact on the endosomal escape of LNP-RNA delivery systems. Compared to lamellar structured LNP-RNA complexes, cubic and inverse hexagonal structured LNP-RNA complexes are able to fuse with endosomal membranes easier, hence leading to higher extent of endosomal escape. These findings provide valuable insight in how nanostructures can affect nanoparticle-cell interactions, but also highlight the potential in utilizing structurally active lipids and nanoparticle structure as an additional handle to controlling the efficacy of drug delivery systems.

## 2 Conclusion

Endosomal escape has been a major hurdle in the development of successful RNA therapeutics. Our studies were able to show that by controlling the nanostructure of LNP-RNA complexes, we can influence the efficiency of endosomal escape without any aid from proteins, endosome acidification, or lipid ionization. Specifically, complexes with cubic structures are able to promote membrane fusion between LNPs and endosomal membranes. This is possible due to their intrinsic ability to lower the elastic cost of the topological transformation from an endosome with a nested LNP to a fused LNP–endosome state having a pore through which RNA can be transported into the cytosol. This enhancement in endosomal escape for cuboplexes led to better RNA delivery compared to its lipoplex counterparts. These results show that nanostructure, and the inclusion of structurally active lipids, is an important factor in designing non-viral delivery systems, and more research is required to fully unravel how nanoparticles of various structures interact with cells.

## 3 Methods

### 3.1 Materials

Glycerol monoooleate (GMO) were purchased from Sigma-Aldrich (MO, USA). 1,2-dioleoyl-*sn*-glycero-3-phosphocholine (DOPC), 1,2-dioleoyl-3-trimethylammonium propane (DOTAP) and 1,2-dioleoyl-*sn*-glycero-3-phosphoethanolamine-N-[methoxy(polyethylene glycol)-2000] (DOPE-PEG) were purchased from Avanti Polar Lipids (AL, USA). siRNA targeting luciferase, siLuc (19 base pair, CUUACGCUGAGUACUUCGA with two 30-deoxythymidine overhangs) was purchased from Dharmacon (Germany). Scrambled siRNA and Alexa Fluor 546 labeled siRNA (AF546-siRNA) were purchased from Qiagen (MD, USA). mRNA targeting firefly luciferase was purchased from Trilink Biotechnologies (CA, USA).

### 3.2 Preparation of LNPs and LNP-RNA complexes

Lipid chloroform solutions were mixed at desired volumetric ratios. Chloroform was removed by treating the solution with a stream of nitrogen and then placing it under vacuum for at least 8 hrs. DOPE-PEG was dried in a separate vial. The dried lipid film was hydrated with sterile Milli-Q water and incubated overnight at 37°C. LNPs were obtained by sonicating the suspension with a cup horn system (Fisher Scientific) for 6 min at 100 % amplitude. The temperature was maintained at below 4 °C during sonication by a water chiller (Qsonica). The LNPs were transferred to the vial with dried DOPE-PEG and incubated for 1 hr at 60 °C for post pegylation.

LNP-RNA complexes were formed by mixing LNPs with siRNA or mRNA at a charge ratio *ρ* (*n*_DOTAP_*/n*_NA_) of 3.

### 3.3 Small angle X-ray scattering

20 mM LNP-RNA complexes were prepared and transferred to quartz capillaries (Hilgenberg Glas, Germany). Synchrotron SAXS were performaed at beamline 12-ID-B of the Advanced Photon Source at Argonne National Laboratory. The average photon energy was 14 keV and the data was radially averaged upon acquisition on a Pilatus 2M detector.

### 3.4 Cell culture

HeLa cells (ATCC), 4T1 cells (ATCC) and HeLa-Luc cells (Signosis) were cultured in full cell media consisting of Dulbecco’s modified Eagle’s medium (Corning), 10% fetal bovine serum (Gibco) and 1% penicillin-streptomycin (Gibco) at 37 °C with 5% of carbon dioxide.

### 3.5 Endosome isolation and characterization

Endosomes were isolated with Trident endosome isolation kit (GeneTex). Particle concentration and size were measured by nanoparticle tracking analysis with Nanosight NS300 (Malvern Panalytical).

### 3.6 Membrane fusion studies

For confocal microscopy analysis, DiD (Biotum) labeled LNPs were prepared by including 0.1% DiD in the lipid mixture. Endosomes were labeled by incubating with 20 *µ*M NileRed for 1 hr at 37 °C. Free dye was removed from endosomes by centrifuging for 30 min at 10,000× g and discarding the supernatant. The endosomes were resuspended in PBS and incubated with DiD labeled LNPs for 6 hrs at 37 °C. The mixture was imaged on an LSM 800 (Carl Zeiss) confocal microscope.

For FRET assay, endosomes were incubated with 20 *µ*M DiO and 20 *µ*M DiI for 1 hr at 37 °C for colabeling. Free dye was removed and endosomes were resuspended in PBS. Endosomes and lipid-siRNA complexes were incubated at 37°C and measured for DiO fluorescence every 5 minutes. Fluorescence was detected with Synergy Neo 2 microplate reader (Biotek).

### 3.7 Live cell imaging of the uptake of LNP-RNA complexes

Labeled LNP-siRNA complexes were prepared by including 0.1% DiI in the lipid mixture. HeLa-Luc were cultured on coverslip bottom dishes (ibidi) and stained with Hoescht dye. Complexes were added to the cells and imaged immediately on an LSM 800 (Carl Zeiss) confocal microscope. 405 nm laser was used for the Hoescht dye channel and 488 nm laser was used for DiO and DiI. Images were acquired every 5 minutes for 3 hrs.

Cells were seeded onto poly-lysine coated glass bottom dishes at 1,000 cells/cm^2^ density, and CellLight™ Early Endosomes-GFP reagent(Thermo Fisher Scientific) was added after cell adherence. After 16 hours of incubation, complexes with Alexa Fluor 546 labeled siRNA were added to the cells with a final siRNA concentration of 33 nM. The dish were observed immediately under confocal microscopy. Images were acquired every 5 minutes for 4 hrs.

### 3.8 Flow cytometry

Cells were seeded onto 12-well plates at 1,000 cells/cm^2^ density prior to transfection. The following day, complexes were added to the cells with a final siRNA concentration of 33 nM, and then incubated for 24 hrs. After incubation, the cells were trypsinized and washed with PBS. Cells were later suspended in FACS buffer (98% PBS, 2% FBS) and analyzed by flow cytometry (BD LSR-Fortessa X-20). After the sample data was acquired, 0.4% Trypan blue was added to the samples at a 1:10 ratio and analyzed by flow cytometry.

### 3.9 Luciferase Assay

Cells were seeded onto 96-well plates at 1,000 cells/cm^2^ density prior to transfection. The following day LNP-mRNA complexes were added to the cells with0.2 *µ*g mRNA per well. The cells were incubated for 24 hrs and the luciferase activity was evaluated by a luciferase assay (Promega). Luminescence was measured with Synergy Neo 2 microplate reader (Biotek), and normalized by the cell protein mass of each well. Cell protein mass was quantified by BCA assay (Thermo Fisher Scientific).

## Supplementary information

Supplementary information is available.

## Acknowledgments

This work was supported by the National Institutes of Health under grant no. 1DP2EB024377-01. This research used resources of the Advanced Photon Source, beamline 12-ID-C, a U.S. Department of Energy (DOE) Office of Science User Facility operated for the DOE Office of Science by Argonne National Laboratory under Contract No. DE-AC02-06CH11357. This research was carried out, in part, in the Materials Research Laboratory Central Research Facilities, University of Illinois.

## Notes

### Competing Interest Statement

The authors have declared no competing interest.

