## Supplementary Information for "Lipid nanoparticle topology regulates endosomal escape and delivery of RNA to the cytoplasm"

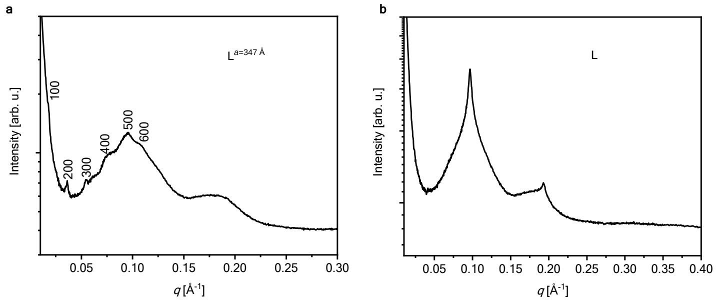


**Fig. 1 Synchrotron SAXS data of LNP-RNA complexes with composition GMO/DOPC/DOTAP/DOPE-PEG.** (**a**) LNP-siRNA complexes were formed with a lipid composition of GMO/DOPC/DOTAP/DOPE-PEG at a molar ratio 25/60/14/1 and charge ratio of 3. The complexes form a lamellar phase with a lattice parameter of 347 Å.(**b**) LNP-mRNA complexes were formed with a lipid composition of GMO/DOPC/DOTAP/DOPE-PEG at a molar ratio 25/60/14/1 and charge ratio of 3. The complexes form a lamellar phase but the exact indexing cannot be determined.

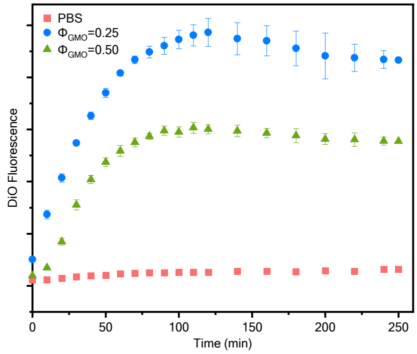


**Fig. 2 FRET assay to evaluate membrane fusion between endosomes and LNP-siRNA complexes with GMO 50% molar percent and GMO 25% molar percent.** The LNP-siRNA complexes with $\Phi_{GMO} = 0.50$ show significant fusion with endosomes. Bouxsein and co-workers [1] showed that siRNA delivered to cell culture by hexagonal bulk phases is efficient but regulated by direct fusion with the plasma membrane compromising its integrity. For that reason, having $Q_{II}$ or $Q_{II}/H_{II}$ is preferable [2] for the purpose of boosting endosomal escape.

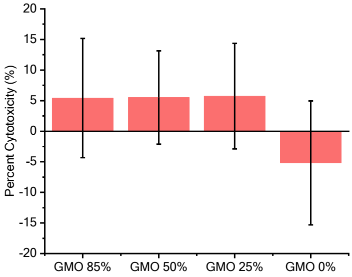


**Fig. 3 Cytotoxicity of LNP-RNA complexes.** Membrane integrity assay of HeLa cells treated with different LNP-RNA complexes.

**Table 1** Sizes of LNP-RNA complexes measured by nanoparticle tracking analysis

| GMO/DOPC/DOTAP/DOPE-PEG | RNA type | Average size (nm) |
|---|---|---|
| 85/0/14/1 | siRNA | 205.2 |
| 50/35/14/1 | siRNA | 190.6 |
| 25/60/14/1 | siRNA | 244.8 |
| 0/85/14/1 | siRNA | 264.9 |
| 85/0/14/1 | mRNA | 152.4 |
| 50/35/14/1 | mRNA | 163.9 |
| 25/60/14/1 | mRNA | 154.5 |
| 0/85/14/1 | mRNA | 200.6 |
